# Physiologic recovery of *Mycobacterium tuberculosis* from drug injury: A molecular study of post antibiotic effect in mice

**DOI:** 10.1101/2025.02.25.640123

**Authors:** Jo Hendrix, Reem Al Mubarak, Adeline Bateman, Lisa M. Massoudi, Karen Rossmassler, Firat Kaya, Matthew D. Zimmerman, Elizabeth A. Wynn, Martin I. Voskuil, Gregory T. Robertson, Camille M. Moore, Nicholas D. Walter

## Abstract

Post-antibiotic effect (PAE) describes the delayed recovery of bacteria following antibiotic exposure. PAE is thought to underlie tuberculosis (TB) treatment forgiveness, *i.e.* the capacity of regimens to tolerate non-adherence. The basis of PAE in *Mycobacterium tuberculosis* (*Mtb*) remains poorly understood, partly because PAE has conventionally been measured based on change in *Mtb* burden *in vitro* rather than change in *Mtb* physiology *in vivo*. We investigated *Mtb* physiologic recovery in the BALB/c mouse model following sub-curative 2- and 4-week durations of the standard isoniazid, rifampin, pyrazinamide, ethambutol (HRZE) treatment. Measurement of rRNA synthesis via the RS ratio® and the entire transcriptome via SEARCH-TB elucidated the dynamics of physiologic recovery.

*Mtb* burden did not increase over 28 days of drug-free post-treatment recovery, indicating prolonged PAE *in vivo.* The RS ratio indicated that *Mtb* ribosomal RNA synthesis resumed within four days of treatment interruption. However, transcriptional changes indicative of metabolic reactivation were delayed for over two weeks. Processes critical for replication, including expression of genes involved in protein and cell wall synthesis, remained suppressed throughout 28 days post-treatment. Longer treatment induced more extensive physiologic perturbation and was associated with slower and less complete recovery.

Expression of processes that are typically induced by environmental stress (*e.g.*, DosR regulon, universal stress proteins, and heat shock proteins) exhibited the reverse, decreasing during drug treatment and rising during recovery.

These findings provide a new basis for understanding PAE based on drug-induced injury and physiologic recovery. Following relatively short durations of HRZE, physiologic recovery of *Mtb* was a slow, sequential and incomplete process *in vivo.* Our observation that longer treatment resulted in even slower recovery suggests that *Mtb* may progressively lose capacity to recover. This work establishes a tractable experimental framework for quantifying the forgiveness of new TB treatment regimens *in vivo*.

## INTRODUCTION

Tuberculosis (TB) remains the leading cause of death from infection worldwide.^1^ There is an urgent need for new treatments that can cure TB more rapidly and reliably.^1^ TB drugs exert post-antibiotic effect (PAE), meaning *Mycobacterium tuberculosis* (*Mtb*) is slow to recover and resume growth following antibiotic exposure.^2,3^ PAE is thought to underly regimen forgiveness, *i.e.* the degree to which a regimen retains efficacy even when patient adherence is sub-optimal.^4^ The World Health Organization has identified forgiveness as a primary criterion for regimen selection.^5^

A decades-old hypothesis is that PAE occurs because bacteria require time to recover from antibiotic-induced damage.^6,7^ Engagement of an antibiotic with its target molecule initiates a secondary injury cascade of drug-specific physiologic perturbations,^8–10^ that can cause macromolecular damage such as DNA breaks^11^ or ribosomal degeneration.^12^ PAE has classically been defined as change in bacterial burden using measures such as colony forming units (CFU),^2,6,13,14^ optical density (OD_600_),^15–17^ or time-to-detection in Bactec culture.^2,18,19^ However, enumeration of bacterial burden provides no information about change in *Mtb* physiology,^20^ the hypothesized basis of PAE. There is a paucity of information about bacterial physiology as *Mtb* recovers during the PAE period.

A second gap in the existing research is that PAE for TB drugs has been evaluated nearly exclusively *in vitro* rather than *in vivo*. For other bacteria, the *in vivo* environment and presence of immunity are known to influence PAE,^21,22^ making PAE substantially longer *in vivo* than *in vitro*. To our knowledge, *in vivo* evaluation of PAE for TB drugs has been limited to a predictive model that suggested that the PAE may be twice as long *in vivo*.^23^ This underscores the need for new methods enabling direct evaluation of PAE on *Mtb in vivo*.

Here, we took a novel approach to evaluating PAE, asking how *Mtb* physiology recovers after drug stress. We used several novel molecular measures to evaluate *Mtb* physiology during the post-antibiotic period after sub-curative treatment durations in the BALB/c high-dose aerosol mouse infection model. In this model, ∼17 weeks of treatment with the standard isoniazid, rifampin, pyrazinamide, and ethambutol (HRZE) regimen is required to achieve non-relapsing cure in 90% of mice.^24^ We interrupted treatment of HRZE after 2 and 4 weeks and sampled during a 28-day post-antibiotic, drug-free recovery period. We used CFU to enumerate bacterial burden and the RS ratio® assay to determine ongoing *Mtb* rRNA synthesis within the mouse lung.^20^ We further used SEARCH-TB, a highly sensitive *Mtb*-targeted RNA-seq platform, to evaluate the entire *Mtb* transcriptome.^8^ These tools enabled us to quantify both the pattern of *Mtb* physiologic injury and adaptation caused by HRZE and the process of recovery after treatment interruption.

Our first goal was to establish the sequence and kinetics by which *Mtb* restores its physiologic processes and resumes growth *in vivo* when drug treatment is interrupted well before cure. Our second goal was to understand if and how the duration of treatment affects *Mtb* recovery. We discovered that while rRNA synthesis increases within 4 days of treatment interruption, the *Mtb* transcriptome changes more slowly, indicating a very gradual recovery. We further found that the longer the duration of drug stress, the more slowly and incompletely *Mtb* recovers during 28 days.

## METHODS

### Murine experiments and RNA extraction

Experiments used the BALB/c high-dose aerosol infection model, which is central to contemporary TB drug development.^13^ Female BALB/c mice, 6 to 8 weeks old, were exposed to aerosol (Glas-Col) with *Mtb* Erdman, resulting in 4.43 log_10_ CFU in lungs on day one. On infection day 11, five mice were euthanized as pre-treatment controls, and HRZE treatment was initiated on the remaining mice via oral gavage five days a week. We used established standard doses of all antibiotics^25^ (Table S1). End of treatment mice (N=4 per treatment duration) were sacrificed on days 12 (2-weeks) and 26 (4-weeks) post-treatment, one day after their final HRZE dose. Additional mice (N=4 per group) were sacrificed on days 4 or 5, 7, 11, 14, 21, and 28 after treatment interruption to assess physiologic recovery (**Fig 1a-b**). Lungs were flash frozen before CFU enumeration and RNA extraction as recently described.^20^ All animal procedures were approved by the Colorado State University Animal Care and Use Committee (Reference number: 1515) and conducted according to established guidelines.

**Fig. 1.**
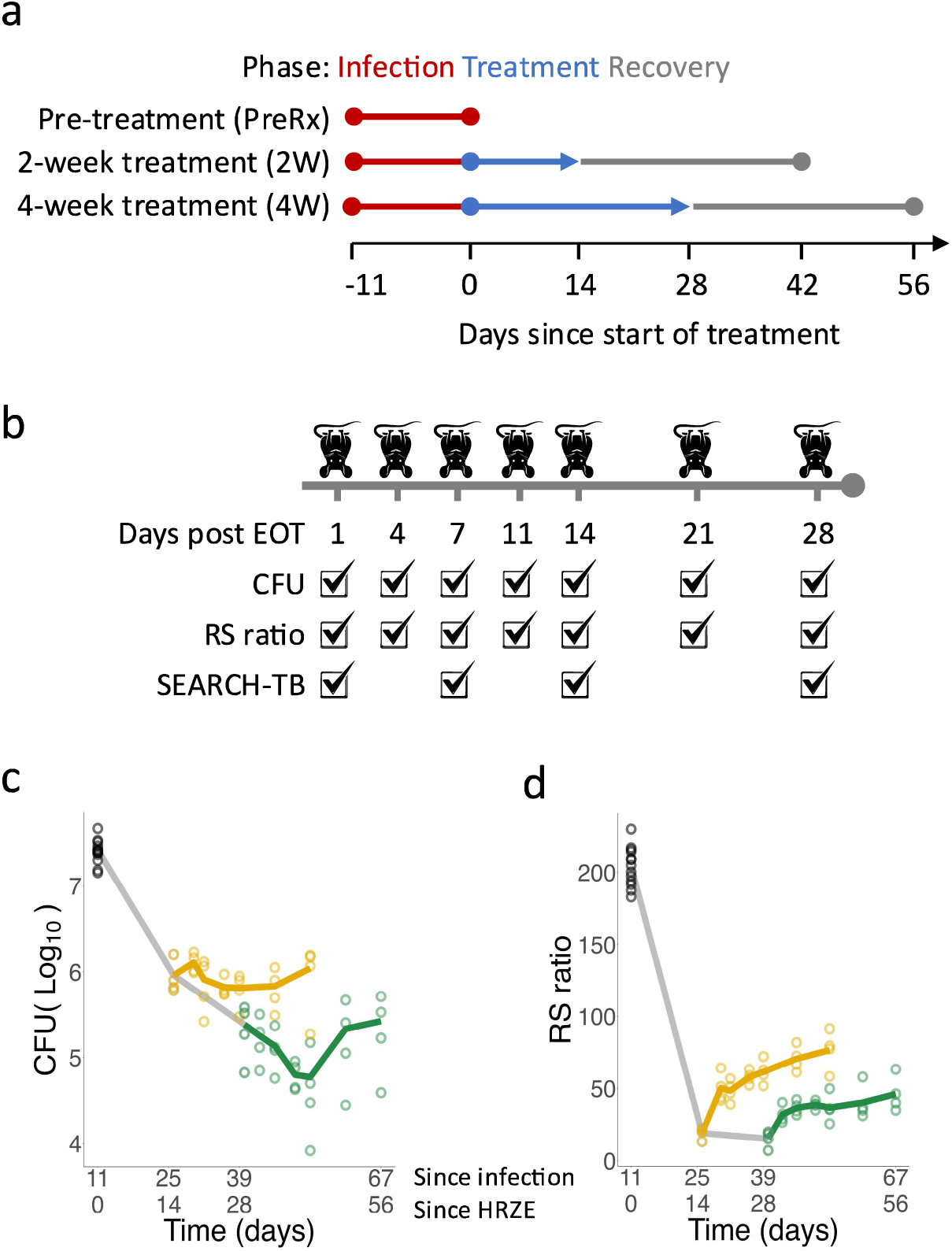
**Schematic of experimental design and eFect of HRZE Treatment**. (a) Timeline of treatment groups. Mice received no treatment for the initial 11 days of infection (red) then were treated for 2 or 4 weeks (blue). The post-treatment recovery phase lasted up to 28 days (gray). (b) Timeline of sample collection during the recovery phase. Five mice were sacrificed at times indicated by mouse icons. Check marks indicate that a given measure (CFU, RS ratio, or SEARCH-TB) was evaluated on a specific day since end of treatment (EOT). (c) CFU burden over time since the start of infection (red dotted line) or start of HRZE treatment (blue dotted line). Each dot represents a mouse from untreated control (black), two-week treatment arm (yellow), or four-week treatment arm (green). A line was drawn through the median values of each time point. (d) RS ratio of *Mtb* displayed as in (a).

### RNA extraction, molecular profiling, and data preparation

RNA extraction, and the RS ratio were performed as previously described.^20,26^ Sequence analysis of samples was performed via SEARCH-TB following recently described methods.^8^ Briefly, RNA was reverse transcribed and cDNA targets were then amplified using the SEARCH-TB panel, following the Illumina AmpliSeq protocol. Libraries were sequenced on an Illumina NovaSeq6000. We followed the bioinformatic analysis and quality control pipeline as described in Supplemental Methods.

### In vitro Minimum Inhibitory Concentration testing

The Minimal Inhibitory Concentration (MIC) was determined for pyrazinamide, rifampicin, ethambutol, or isoniazid against Mtb Erdman in 7H9 media supplemented with 0.2% [v:v] glycerol and 10% [v:v] ADC, with 0.05% [v:v] Tween-80 (7H9 media), pH 6.6. MIC values were determined by a broth microdilution assay using two-fold serial drug dilutions. The lowest consecutive antimicrobial concentration that showed a ≥ 80% reduction in OD_600_ relative to drug-free control wells, was regarded as the MIC.

### Pharmacokinetic measurements

After euthanizing mice individually, blood was collected via cardiac puncture into K3EDTA tubes (Greiner Bio-One MiniCollect 1 mL K3E K3EDTA item#450474), maintained on ice before centrifugation at 10,000 RCF for 2 minutes at 4°C, plasma was transferred, then frozen at-80°C within 1 hour of collection. Drug levels in plasma were quantified by high pressure liquid chromatography coupled to tandem mass spectrometry (LC-MS/MS) using previously documented methods for isoniazid,^27^ rifampicin,^27^ pyrazinamide,^28^ and ethambutol.^29^

### Transcriptome comparisons

We evaluated the effect of HRZE treatment by comparing gene expression at the end of 2-or 4-week treatment with pre-treatment controls. We evaluated recovery by comparing: (1) the end of 28-day recovery with the end of 2-or 4-week treatment, (2) the end of 28-day recovery with pre-treatment controls and (3) the end of 28-day recovery following 4-week treatment with the end of 28-day recovery following 2-week treatment. For each gene, negative binomial generalized linear models were fit using edgeR^30,31^ to identify the effect of treatment and recovery. For each comparison of interest, likelihood ratio tests were performed and genes with Benjamini–Hochberg adjusted P-value^32^ less than 0.05 were considered significantly differentially expressed.

### Bacterial Average Expression Comparison

The average normalized gene expression for genes in categories established by Cole et al.^33^ or curated from the literature (**Table S2**) were calculated for each sample as previously described.^8^ Values were compared between pairs of timepoints using a t-test.

Gene categories with Benjamini–Hochberg adjusted P-values^34^ less than 0.05 were considered significant. Differential expression, functional enrichment, and visualizations can be evaluated interactively using an Online Analysis Tool created using the R package Shiny^35^ v1.8.1. [https://microbialmetrics.org/analysis-tools/]

## RESULTS

### EFect of HRZE treatment on *Mtb* burden and physiologic state

The median CFU burden in pre-treatment control mice was 7.4 log. HRZE rapidly reduced the burden of culturable *Mtb* in mouse lungs, decreasing CFU 97.2% (1.6 log) and 99.0% (2.0 log) after two and four weeks, respectively (**Fig 1c**). Among the pre-treatment control mice, the median RS ratio was 204, indicating a high rate of rRNA synthesis prior to drug exposure. Two and four weeks of HRZE reduced the median RS ratio to 20.5 and 17.0 (**Fig 1d**), respectively, indicating that drug stress slowed rRNA synthesis.

Treatment with HRZE for two and four weeks massively transformed the *Mtb* transcriptome, significantly altering expression of 2,497 (70.3%) and 2,617 (73.7%) genes relative to pre-treatment control, respectively (Supplemental **Fig S1**). The expression changes closely reproduced HRZE-induced changes we previously reported in a separate experiment^8^ (Supplemental **Fig S2**). As observed previously, the 4-week transcriptome was a somewhat more extreme version of the 2-week transcriptome (Supplemental **Fig S3**). The number of genes differentially expressed in direct comparison between 2 and 4 weeks was modest (N=229), but the average absolute log_2_ fold change of significant genes was larger at 4-weeks (1.48) than at 2 weeks (1.35). Treatment-induced changes in specific physiologic categories are discussed below and can be explored interactively via our Online Analysis Tool (https://microbialmetrics.org/analysis-tools/).

### PK during the recovery period

In the 4-week treatment group, we tested plasma drug concentrations one day after the final treatment dose. Pyrazinamide was detected at a trace (8.1ng/mL) concentration [inhibitory concentration reported as 6,000 to 50,000 ng/mL at pH 5.5^36^, and > 64,000 ng/mL at pH 6.6, as tested here]. Ethambutol was detected at a sub-inhibitory (31.5ng/mL) concentration [inhibitory concentration = 1,000 ng/mL]. Rifampin was detected at an inhibitory (565.5ng/mL) concentration [inhibitory concentration = 8 ng/mL]. We additionally tested plasma drug concentrations five days after the final treatment dose and measured levels were BLQ for of rifampin (LOQ-1ng/mL) and pyrazinamide (LOQ-25ng/mL). One of four mice tested had trace (1.1 ng/mL) concentration of isoniazid [inhibitory concentration = 30 ng/mL]. All four mice tested had trace levels of ethambutol (2.49 ng/mL).

### Change in *Mtb* burden during the post-antibiotic recovery period

When HRZE was stopped after 2 or 4 weeks, CFU counts stabilized. CFU did not increase significantly during the subsequent 28-day post-antibiotic recovery period (**Fig 1c**) (Supplemental **Table S3-4**).

### Change in *Mtb* rRNA synthesis during the post-antibiotic recovery period

When HRZE was stopped after 2 weeks, the median RS ratio rebounded from 20.5 at end of treatment to 47.5 four days later, indicating a greater than 2-fold increase in rRNA synthesis. (**Fig 1d**). Thereafter, the RS ratio increased more gradually, reaching 78.6 at the end of the 28-day post-antibiotic recovery period, a level characteristic of chronic immune-contained infection.^20^ When HRZE was stopped after 4 weeks, the RS ratio recovered more gradually, with significantly lower values than in the 2-week group at all timepoints (least significant Adj-P=0.049). (Supplemental **Table S5-6)**

### Change in *Mtb* transcriptome during the post-antibiotic recovery period

The *Mtb* transcriptome recovered slowly (**Fig 2a**). Following 2-week treatment, the number of genes differentially expressed relative to end of treatment was 32, 150 and 800 on days 7, 14 and 28 post-treatment, respectively (**Fig 2a**). Following 4-week treatment, transcriptional change lagged behind the 2-week treatment with 16, 38 and 731 genes differentially expressed on days 7, 14 and 28, respectively (**Fig 2b**). For both the 2-and 4-week groups, this slow pace of transcriptional change is reflected in the PCA plots, where samples from days 7 and 14 post-treatment were similar to the end of treatment samples, while day 28 was distinct along PC1 (**Fig 2c-d**).

**Fig. 2.**
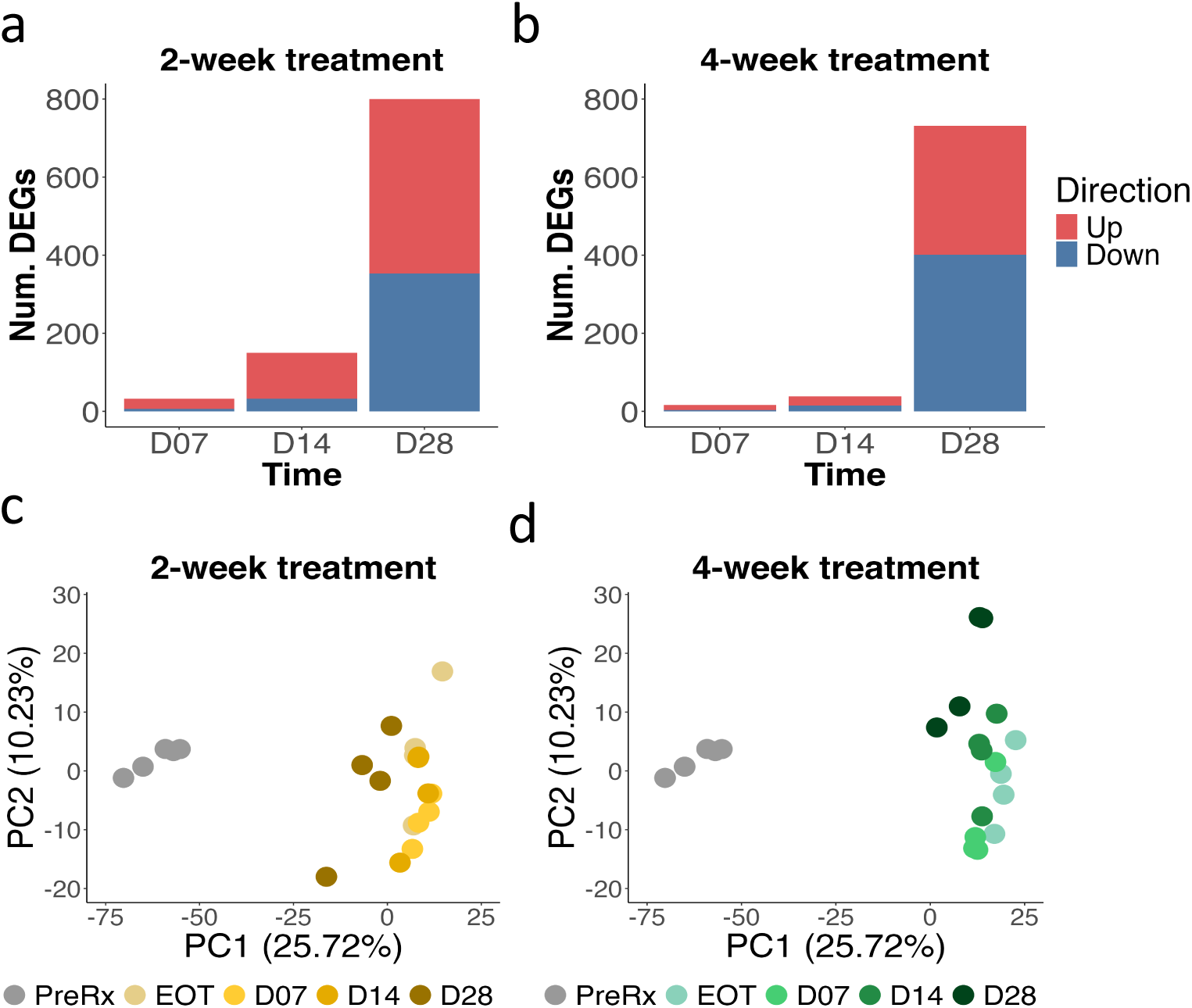
Transcriptional recovery dynamics. (a-b) Number of differentially expressed genes at recovery day 7, 14, and 28 compared to end of treatment after two (a) or four (b) weeks of treatment. Genes significantly down-and up-regulated relative to control (adj. P < 0.05) are shown in blue and red, respectively. (c-d) PCA plots of pre-treatment (gray), end of treatment and recovery time points for two (c) and four (d) week treatment groups.

### Recovery of specific *Mtb* physiologic processes

#### Central carbon metabolism related genes: Suppression during treatment and rebound during recovery

During treatment, decreased expression of the following gene sets indicated a global decrease in central carbon metabolism relative to pre-treatment control: (1) glycolysis related genes (Adj-*P*<0.0001 and 0.00042 after 2-and 4-weeks, respectively), (2) pentose phosphate pathway (Adj-*P*=0.00033 and 0.00059 after 2-and 4-weeks, respectively), (3) pyruvate dehydrogenase genes (Adj-*P*<0.0001 and 0.00074 after 2-and 4-weeks, respectively), (4) beta-oxidation genes (Adj-*P*<0.0001 and 0.00095 after 2-and 4-weeks, respectively), (5) glyoxylate bypass genes (Adj-*P*=0.0012 and <0.0001after 2-and 4-weeks, respectively), (6) cholesterol side chain degradation (Adj-*P*<0.0001 and 0.00012 after 2-and 4-weeks, respectively), (7) cholesterol A and B ring degradation (Adj-*P*<0.0001 and 0.00016 after 2-and 4-weeks, respectively) and (8) cholesterol C and D ring degradation (Adj-*P*=0.031 and 0.018 after 2-and 4-weeks, respectively) (**Fig 3a**).

**Fig. 3.**
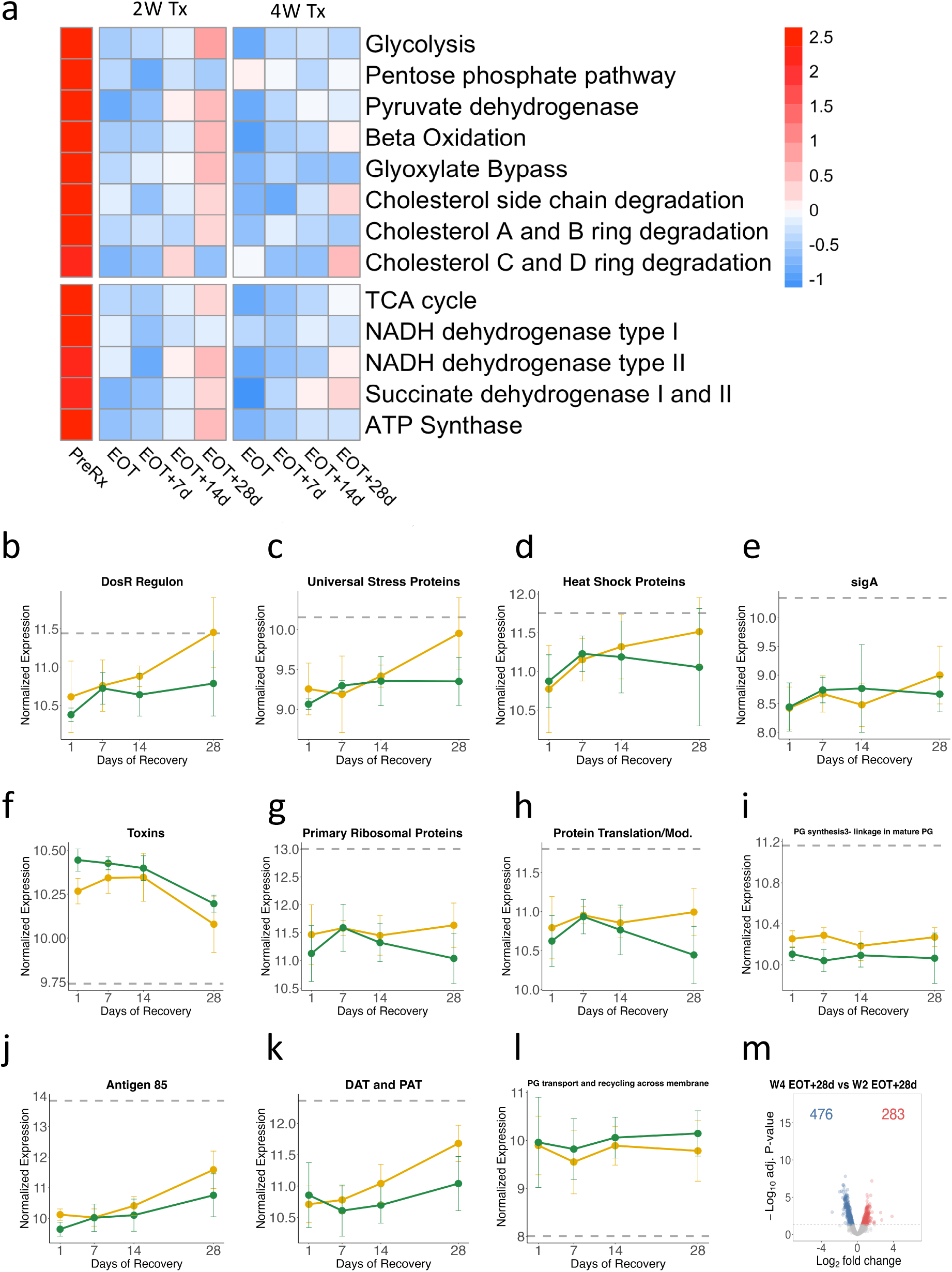
Transcriptional recovery of specific *Mtb* gene categories. (a) Normalized gene expression of select gene categories involved in central carbon metabolism (top) and electron transport (bottom) for the pre-treatment control, end of two and four weeks of treatment, and after 7, 14, and 28 days of recovery. (b-l) Normalized gene expression of select gene categories during treatment interruption following two (yellow) or four (green) weeks of treatment. Points indicate the average expression level and whiskers indicate 95% Confidence Interval between the five samples per time point. Gray line indicates average expression level of pre-treatment control. (m) Volcano plot showing log_2_ fold changes and −log_10_ P-values of genes differentially expressed after 28 days of treatment interruption following four weeks of treatment compared to two weeks of treatment. Genes significantly down-and up-regulated relative to control (adj. P < 0.05) are shown in blue and red, respectively.

Conversely, during recovery, expression of central-carbon related genes gradually increased relative to end of treatment, albeit generally to a greater degree after 2-week treatment than after 4-week treatment. For example, expression of glycolysis genes increased significantly during recovery from 2-week treatment (Adj-P=0.039) but not from 4-week treatment (**Fig 3a**). Line plots for each gene set discussed above are included in Supplemental Information (Supplemental **Fig S4**).

#### Electron transport related genes: Suppression during treatment and rebound during recovery

During treatment, expression of the following electron transport-related genes decreased significantly relative to pre-treatment control: (1) TCA cycle genes (Adj-*P*<0.0001 for 2-and 4-weeks), (2) NADH dehydrogenase type I genes (Adj-*P*=0.0020 for 2-and 4-week treatment), (3) NADH dehydrogenase type II genes (Adj-*P*=0.011 and 0.0019 after 2-and 4-weeks, respectively), (4) succinate dehydrogenase types I and II (Adj-*P*=0.00031 and <0.0001 after 2-and 4-weeks, respectively) and (5) ATP synthetase genes (Adj-*P*=0.00014 and 0.0016 after 2-and 4-weeks, respectively) (**Fig 3a**).

Conversely, during recovery, expression of respiration related genes gradually increased relative to end of treatment, albeit to a greater degree after 2-week treatment than after 4-week treatment. An example is expression of ATP synthetase genes which increased significantly after 28 days of recovery for the 2-week treatment (Adj-P=0.032) but not for the 4-week treatment (**Fig 3a**). At the end of the 28-day post-antibiotic recovery phase, expression of ATP synthase genes remained lower for the 4-week treatment group compared to the 2-week group (**Fig 3a**).

During treatment, oxidative phosphorylation appeared to transition from the primary cytochrome *bcc/aa3* supercomplex (significantly decreased expression after treatment [Adj-P<0.0001 for 2-and 4-week treatment]) (**Fig 3a**) to the less-e=icient alternative cytochrome *bd* oxidase (increased expression at weeks 2 [Adj-P=0.064] and 4 [Adj-P=0.010]) (**Fig 3a**). This was partially reversed with significantly increased expression of cytochrome bcc/aa3 genes after 28 days of recovery from the 2-week treatment (Adj-P=0.034) but not after recovery from the 4-week treatment. Expression of cytochrome bd did not decrease during the recovery period. Line plots for each gene set discussed above are included in Supplemental Information (Supplemental **Fig S5**).

#### DosR regulon genes: Suppression during treatment and rebound during recovery

During treatment with HRZE, expression of the DosR regulon, that responds to exposure to hypoxia, nitric oxide (NO), and carbon monoxide,^37–39^decreased significantly relative to pre-treatment control (Adj-*P <*0.0093 and 0.0033 after 2-and 4-weeks, respectively). Conversely, during recovery after 2-week treatment, DosR expression gradually rose (Adj-P=0.046) in parallel with the resumption of aerobic metabolism. During recovery after 4-week treatment, DosR expression rose more slowly than after 2-week treatment, concordant with the slower metabolic recovery observed after 4-week treatment. At the end of the 28-day post-antibiotic recovery phase, DosR expression remained lower for the 4-week treatment group compared to the 2-week group, though not significantly (Adj-*P=*0.12) (**Fig 3b**).

Six DosR regulon genes plus one additional gene are described as universal stress proteins (USP) because they are induced by diverse environmental stresses.^40^ During treatment, USP decreased significantly relative to pre-treatment control (Adj-*P*=0.0030 and 0.0027 after 2-and 4-weeks, respectively). Conversely, during recovery after 2-week treatment, expression of USP genes rose, but did not reach statistical significance (Adj-*P*=0.055) (**Fig 3c**). During recovery after 4-week treatment, USP expression rose more slowly than after 2-week treatment.

#### Heat shock genes: Suppression during treatment and rebound during recovery

During treatment, expression of genes for heat shock proteins (HSP) that serve as molecular chaperones in protein folding and are induced by various environmental stresses that disrupt protein folding^41^ decreased significantly relative to pre-treatment control (Adj-*P*=0.014 and 0.0031 after 2-and 4-weeks, respectively). During recovery after 2-week treatment, expression of HSP genes rose more than after 4-week treatment (**Fig 3d**).

#### Change in sigma factors

During treatment, expression of the gene for the growth-enabling primary sigma factor SigA, was profoundly suppressed relative to pre-treatment control (Adj-*P*=2.9×10^-17^ for both 2-and 4-week treatments). Conversely, during recovery after 2-week treatment, expression of *sigA* rose modestly but significantly (Adj-*P*=0.046). During recovery after 4-week treatment, expression of *sigA* did not change significantly (Adj-*P=*0.55). Other sigma factors were variably expressed during treatment and recovery (**Fig 3e**).

#### Toxin genes: Induction during treatment and resolution during recovery

During treatment, expression of toxins, post-transcriptional stress responses which inhibit replication, translation, and cell wall synthesis, and metabolism,^42^ increased significantly relative to pre-treatment control (Adj-*P*=0.00042 and 0.00016 after 2-and 4-weeks, respectively). Conversely, during recovery, expression of toxins decreased though only significantly for 4-weeks (Adj-*P*=0.017) (**Fig 3f**).

#### Protein synthesis-related genes: Suppression during treatment and lack of change during recovery

During treatment, expression of primary ribosomal protein genes (*i.e.,* after exclusion of C-alternative ribosomal protein genes) decreased significantly relative to pre-treatment control (Adj-*P*=0.0039 and 0.0018 after 2-and 4-weeks, respectively). Similarly, during treatment, expression of genes associated with protein translation and modification decreased significantly (Adj-*P*=0.0043 and 0.0013 after 2-and 4-weeks, respectively).

During the 28-day recovery period, neither the 2-week nor 4-week treatment group displayed significant increase in expression of genes for primary ribosomal proteins or protein translation and modification, suggesting minimal recovery of protein synthesis (**Fig 3g-h**).

#### Change in cell wall-associated genes

Expression of genes related to *de novo* synthesis of cell wall constituents decreased during treatment and did not rise during recovery. Specifically, during treatment, expression of the following genes related to mycolic acid synthesis decreased significantly relative to pre-treatment control: (1) *fas,* the gene coding for fatty acid synthase I, the first step mycolic acid synthesis (Adj-*P*=0.013 and 2.2×10^-5^ after 2-and 4-weeks, respectively) (2) fatty acid synthase II (FAS-II) genes (Adj-*P*=0.00047 and 0.0010 after 2-and 4-weeks, respectively), (3) mycolic acid modification genes (Adj-*P*=0.0021 and <0.0001 after 2-and 4-weeks, respectively) and (4) mycolic acid transfer and modification genes (Adj-*P*=0.00068 and 0.00058 after 2-and 4-weeks, respectively). During the 28-day recovery period after either the 2-or 4-week treatment, there was no significant increase in expression of these mycolic acid synthesis genes. Similarly, expression of genes for phthiocerol dimycocerosate (PDIM) decreased significantly during treatment (Adj-*P*=0.0059 and 0.0011 after 2-and 4-weeks, respectively) and did not rise significantly during 28-day recovery. Finally, expression of genes involved in linking peptidoglycan for cell wall synthesis decreased significantly during treatment (Adj-*P*<0.0001 for 2-and 4-weeks) and did not rise significantly during 28-day recovery (**Fig 3i**).

Despite the evidence above that *de novo* synthesis of cell wall constituents did not increase after treatment interruption, there was evidence of cell wall remodeling and recycling during recovery. Specifically, expression of genes for Antigen 85 (Ag85), a mycolyl transferase that has a role in cell wall remodeling, decreased significantly during treatment relative to pre-treatment control (Adj-*P*<0.0001 for 2-and 4-weeks). Conversely, during recovery, expression of Ag85 genes rose significantly for the 2-week treatment (Adj-*P*=0.034) but not 4-week treatment (**Fig 3j**). Expression of genes related to synthesis of trehalose containing glycolipids diacyltrehalose (DAT) and pentaacyltrehalose (PAT) decreased significantly during treatment (Adj-*P*=4.6E-5 and 0.0017 after 2-and 4-weeks, respectively). Conversely, during recovery, expression of DAT and PAT genes rose significantly for the 2-week (Adj-*P*=0.00024) but not 4-week treatment group (**Fig 3k**).

Finally, expression of the uspA,B,C regulon (ABC transporters) involved in peptidoglycan recycling across the membrane increased significantly during treatment (Adj-P=0.0014 and 0.0033 after 2-and 4-weeks, respectively) (**Fig 3l**).

### Transcriptional recovery was incomplete

At the end of the 28-day drug-free recovery period, the *Mtb* transcriptome remained markedly altered with 2,294 (64.6%) genes differentially expressed in the 2-week treatment group and 2,546 (71.7%) in the 4-week treatment group relative to pre-treatment control, respectively. Persistent differences between the end of recovery *Mtb* phenotypes and pre-treatment are also highlighted in PCA (**Fig 2c-d**).

### Sustained diFerences between 2-and 4-week treatments

During the recovery period, the transcriptional patterns of the 2-and 4-week treatment groups grew more different rather than more similar. Specifically, at the end of treatment, there were 229 genes differentially expressed between 2-and 4-week treatments. After the 28-day recovery period, the number of genes differentially expressed between 2-and 4-week treatment groups rose to 759 (Supplemental Fig S6) (**Fig 3m**).

## DISCUSSION

By applying several novel molecular methods to evaluate PAE we elucidated the physiological recovery of *Mtb* in the BALB/c mouse during the recovery period after sub-curative treatment durations with HRZE. We discovered that recovery from the quiescent drug-injured end of treatment *Mtb* phenotype was slow and sequential with gradual reactivation of metabolism and alteration in canonical stress responses. However, expression of genes associated with protein synthesis and *de novo* synthesis of cell wall constituents remained unchanged even 28 days after treatment interruption, suggesting a prolonged delay in reinitiation of replication. *Mtb* recovery depended on the duration of antibiotic exposure; longer treatment appeared to induce a more extensive pattern of injury and adaptation that was associated with slower physiologic recovery. By focusing on recovery of physiologic processes rather than bacterial burden, this molecular assessment expands understanding of PAE and establishes an experimental model with potential to quantify TB regimen forgiveness *in vivo*.

PAE is the continued suppression of bacterial growth observed after transient antibiotic exposure (*i.e.,* beyond the point at which drug or drugs are entirely cleared).^43^ In this study, all drugs were undetectable on the fifth day after treatment (with the exception of a single mouse with trace detection of isoniazid), providing confidence that our results indicate PAE rather than lingering residual drug exposure. Engagement of drugs with their molecular targets is known to initiate a secondary cascade of injury and adaptation^8,9,44,45^ that may damage key cellular processes^6,12^ and deplete essential metabolites.^3^ PAE is thought to represent the repair and repletion period required before growth resumes; however, understanding has been limited by the methods conventionally used to measure PAE. PAE is generally quantified *in vitro* based on the length of the lag before there is a rise in bacterial burden,^46^ a measure that provides no insight into change in *Mtb* physiology.

Additionally, PAE *in vitro* may fail to recapitulate PAE *in vivo* where bacterial recovery may be modulated by host immunity.^23^ As our results illustrate, CFU did not rise discernably during 28 days after treatment interruption, making it effectively impossible to measure PAE based on change in *Mtb* burden in the BALB/c mouse model.

Here, we assessed PAE *in vivo* in a new way, using molecular assays to interrogate bacterial physiology rather than bacterial burden. Our first key finding was that gradual physiologic recovery was measurable even though CFU remained unchanged. After a lag, transcriptional change between post-treatment days 14 and 28 indicated a reboot of metabolism, remodeling of the cell wall, and regulatory changes. Equally notable were processes that did not change during the post-treatment recovery period which included growth-associated processes such as genes coding for synthesis of primary ribosomal proteins and cell wall constituents. The lack of change in genes required for replication is consistent with the lack of change in CFU during the recovery period and suggests that recovery during a 28-day period was limited to repair rather than replication.

Second, our findings point to important differences between *Mtb* responses to drug exposure and to environmental stressors.^47^ For instance, the DosR regulon was initially called the dormancy regulator since it is induced by environmental conditions that inhibit aerobic respiration and arrest growth. However, in both our previous work^8^ and this project, we identified the opposite association: as HRZE treatment shifted *Mtb* to a quiescent phenotype, expression of the DosR regulon decreased rather than increased. After drug stress was removed and *Mtb* slowly recovered, expression of DosR increased in parallel with increased expression of *Mtb* metabolism-related genes. An additional example of divergence between responses to environment stress and drugs is HSPs that are induced by various environmental stresses.^41^ We found that expression of HSPs was suppressed during drug stress and increased during post-treatment recovery. In the aggregate, these findings highlight that responses to and recovery from drug treatment are distinct from responses to environmental stress.

A third key finding was that both SEARCH-TB and RS ratio suggested that *Mtb* had impaired recovery from 4-week drug stress, relative to 2-week drug stress. Nearly all processes that changed during the recovery period changed more slowly or to a lesser degree after 4-week treatment than after 2-week treatment. Even 28 days after treatment interruption, *Mtb* treated for four weeks remained transcriptionally distinct from *Mtb* treated for two weeks. An association between duration of drug exposure and PAE was noted during the earliest studies with streptococcus^48^ and subsequently shown with other bacteria,^49^ but has not to our knowledge been shown for *Mtb* either *in vitro* or *in vivo*. Our observation that both the degree of transcriptional injury and impairment of recovery increase with longer treatment (*i.e.,* four versus two weeks) suggests that *Mtb* may become progressively depleted and less capable of recovery over time. It is conceivable that the cascade of drug-induced injury leads to a tipping point at which homeostasis can no longer be sustained, resulting in “death by depletion.”

Finally, we found that even 28 days post-treatment interruption, *Mtb* recovered incompletely, continuing to have a more quiescent phenotype than that observed in pre-treatment control mice. The RS ratio remained <60% of the pre-treatment level and expression of genes indicative of *Mtb* replication continued to be expressed at lower levels than pre-treatment. A potential explanation is change in the host immune milieu. The pre-treatment control mice were sacrificed on infection day 11, prior to the onset of adaptive immunity which occurs 16-20 days post-infection in the BALB/c mouse.^50^ We have previously shown that the onset of adaptive immunity in the BALB/c mouse is associated with slowing of *Mtb* rRNA synthesis, suggesting immune containment results in a more quiescent *Mtb* phenotype.^20^ The continued quiescent phenotype observed at 28 days of treatment interruption may not solely represent residual antibiotic damage, but also indicate more effective immune containment.

The clinical concept of regimen forgiveness (capacity to tolerate non-adherence) is thought to be a function of both pharmacokinetics (*i.e.,* lingering subinhibitory drug concentrations) and PAE.^5,51^ Here we observed that, weeks after drugs have been cleared, *Mtb* physiologic recovery from drug-induced injury is slow. The observation in human patients that missing doses in a clustered fashion increases risk of treatment failure more than missing doses randomly seems consistent with the idea that *Mtb* requires time for physiologic recovery.^52^

This study had several limitations. We selected 28 days as our longest post-antibiotic timepoint with the expectation that *Mtb* would have fully recovered from drug exposure. It is conceivable that even longer post-treatment observations might have revealed further long-term transcriptional *Mtb* recovery. Second, here we tested only the standard regimen. Because individual drugs induce different patterns of physiologic injury and adaptation *in vivo*^8^ and have variable PAE *in vitro*,^2,18,49,53^ it is plausible that different combination regimens will have distinct PAE. Molecular evaluation of additional regimens will be a next step. Finally, as described above, our findings were likely influenced by the onset of adaptive immunity. Repeating this study in an immunodeficient mouse model could isolate antibiotic recovery from the effects of the immune system.

This study reevaluated PAE by elucidating *Mtb* physiological recovery after sub-curative treatment with the standard HRZE regimen. By employing molecular assays of physiologic processes rather than traditional measures of bacterial burden, we probed physiologic injury during treatment and recovery that are the putative basis of PAE. This revealed that *Mtb* recovery *in vivo* is a slow sequential process and that the longer the treatment, the slower the recovery. Future investigations into the influence of specific drug combinations, longer recovery periods, and immune interactions will further clarify the mechanisms underlying PAE and could inform the development of more effective TB treatment strategies that capitalize on the physiological constraints of *Mtb*.

## REFERENCES

1. WHO. Global tuberculosis report 2024. https://www.who.int/publications/i/item/9789240101531.

2. Chan, C.-Y., Au-Yeang, C., Yew, W.-W., Hui, M. & Cheng, A. F. B. Postantibiotic Effects of Antituberculosis Agents Alone and in Combination. Antimicrob. Agents Chemother. 45, 3631–3634 (2001).

3. MacKenzie, F. M. & Gould, I. M. The post-antibiotic effect. J. Antimicrob. Chemother. 32, 519–537 (1993).

4. Susanto, B. O., Wicha, S. G., Hu, Y., Coates, A. R. M. & Simonsson, U. S. H. Translational Model-Informed Approach for Selection of Tuberculosis Drug Combination Regimens in Early Clinical Development. Clin. Pharmacol. Ther. 108, 274–286 (2020).

5. WHO. Target regimen profiles for tuberculosis treatment, 2023 update. https://www.who.int/publications/i/item/9789240081512.

6. Eagle, H., Fleischman, R. & Musselman, A. D. The bactericidal action of penicillin in vivo: the participation of the host, and the slow recovery of the surviving organisms. Ann. Intern. Med. 33, 544–571 (1950).

7. Bigger, J. W. The bactericidal action of penicillin on Staphylococcus pyogenes. Ir. J. Med. Sci. 1926*-*1967 19, 553–568 (1944).

8. Wynn, E. A. et al. Transcriptional adaptation of Mycobacterium tuberculosis that survives prolonged multi-drug treatment in mice. mBio 14, e02363–23 (2023).

9. Koul, A. et al. Delayed bactericidal response of Mycobacterium tuberculosis to bedaquiline involves remodelling of bacterial metabolism. Nat. Commun. 5, 3369 (2014).

10. Poonawala, H. et al. Transcriptomic responses to antibiotic exposure in Mycobacterium tuberculosis. Antimicrob. Agents Chemother. 68, e01185–23 (2024).

11. Saito, K. Oxidative damage and delayed replication allow viable Mycobacterium tuberculosis to go undetected. https://www.science.org/doi/10.1126/scitranslmed.abg2612 (2021) doi:10.1126/scitranslmed.abg2612.

12. Maiväli, Ü., Paier, A. & Tenson, T. When stable RNA becomes unstable: the degradation of ribosomes in bacteria and beyond. Biol. Chem. 394, 845–855 (2013).

13. Gumbo, T., Lenaerts, A. J., Hanna, D., Romero, K. & Nuermberger, E. Nonclinical Models for Antituberculosis Drug Development: A Landscape Analysis. J. Infect. Dis. 211, S83– S95 (2015).

14. Alotaibi, S. H. Study of drug interaction, mutant frequency and mutant prevention concentration of DFMBT against *Mycobacterium tuberculosis* H37RV. Microb. Pathog. 176, 106023 (2023).

15. Hurdle, J. G. et al. A microbiological assessment of novel nitrofuranylamides as anti-tuberculosis agents. J. Antimicrob. Chemother. 62, 1037–1045 (2008).

16. Yu, X. et al. Nosiheptide Harbors Potent In Vitro and Intracellular Inhbitory Activities against Mycobacterium tuberculosis. Microbiol. Spectr. 10, e01444–22.

17. Islam, M. I. et al. Antimicrobial activity of IDD-B40 against drug-resistant Mycobacterium tuberculosis. Sci. Rep. 11, 740 (2021).

18. Chan, C.-Y., Au-Yeang, C., Yew, W.-W., Leung, C.-C. & Cheng, A. F. B. In Vitro Postantibiotic Effects of Rifapentine, Isoniazid, and Moxifloxacin against Mycobacterium tuberculosis. Antimicrob. Agents Chemother. 48, 340–343 (2004).

19. Reddy, V. M., Einck, L., Andries, K. & Nacy, C. A. In Vitro Interactions between New Antitubercular Drug Candidates SQ109 and TMC207. Antimicrob. Agents Chemother. 54, 2840–2846 (2010).

20. Walter, N. D. et al. Mycobacterium tuberculosis precursor rRNA as a measure of treatment-shortening activity of drugs and regimens. Nat. Commun. 12, 2899 (2021).

21. Craig, W. A. Post-antibiotic effects in experimental infection models: relationship to in-vitro phenomena and to treatment of infections in man. J. Antimicrob. Chemother. 31, 149–158 (1993).

22. Craig, W. A. The postantibiotic effect. Clin. Microbiol. Newsl. 13, 121–124 (1991).

23. Wagh, S. et al. Model-Based Exposure-Response Assessment for Spectinamide 1810 in a Mouse Model of Tuberculosis. Antimicrob. Agents Chemother. 65, 10.1128/aac.01744-20 (2021).

24. Berg, A. et al. Model-Based Meta-Analysis of Relapsing Mouse Model Studies from the Critical Path to Tuberculosis Drug Regimens Initiative Database. Antimicrob. Agents Chemother. 66, e01793–21.

25. Chen, C. et al. Assessing Pharmacodynamic Interactions in Mice Using the Multistate Tuberculosis Pharmacometric and General Pharmacodynamic Interaction Models. CPT Pharmacomet. Syst. Pharmacol. 6, 787–797 (2017).

26. Walter, N. D. et al. Lung microenvironments harbor Mycobacterium tuberculosis phenotypes with distinct treatment responses. Antimicrob. Agents Chemother. 67, e00284–23.

27. Xu, Y. et al. Matrix metalloproteinase inhibitors enhance the e=icacy of frontline drugs against Mycobacterium tuberculosis. PLoS Pathog. 14, e1006974 (2018).

28. Irwin, S. M. et al. Bedaquiline and Pyrazinamide Treatment Responses Are Affected by Pulmonary Lesion Heterogeneity in Mycobacterium tuberculosis Infected C3HeB/FeJ Mice. ACS Infect. Dis. 2, 251–267 (2016).

29. Zimmerman, M. et al. Ethambutol Partitioning in Tuberculous Pulmonary Lesions Explains Its Clinical E=icacy. Antimicrob. Agents Chemother. 61, e00924–17 (2017).

30. Robinson, M. D., McCarthy, D. J. & Smyth, G. K. edgeR: a Bioconductor package for differential expression analysis of digital gene expression data. Bioinforma. Oxf. Engl. 26, 139–140 (2010).

31. McCarthy, D. J., Chen, Y. & Smyth, G. K. Differential expression analysis of multifactor RNA-Seq experiments with respect to biological variation. Nucleic Acids Res. 40, 4288– 4297 (2012).

32. Benjamini, Y. & Hochberg, Y. Controlling the False Discovery Rate: A Practical and Powerful Approach to Multiple Testing. J. R. Stat. Soc. Ser. B Methodol. 57, 289–300 (1995).

33. Cole, S. T. et al. Deciphering the biology of Mycobacterium tuberculosis from the complete genome sequence. Nature 396, 190–190 (1998).

34. Murtagh, F. & Legendre, P. Ward’s Hierarchical Agglomerative Clustering Method: Which Algorithms Implement Ward’s Criterion? J. Classif. 31, 274–295 (2014).

35. Chang, W., et al. shiny: Web Application Framework for R. (2024).

36. Handbook of anti-tuberculosis agents. Introduction. Tuberc. Edinb. Scotl. 88, 85–86 (2008).

37. Voskuil, M. I. et al. Inhibition of Respiration by Nitric Oxide Induces a Mycobacterium tuberculosis Dormancy Program. J. Exp. Med. 198, 705–713 (2003).

38. Voskuil, M. I., Bartek, I., Visconti, K. & Schoolnik, G. K. The Response of Mycobacterium Tuberculosis to Reactive Oxygen and Nitrogen Species. Front. Microbiol. 2, (2011).

39. Shiloh, M. U., Manzanillo, P. & Cox, J. S. Mycobacterium tuberculosis Senses Host-Derived Carbon Monoxide during Macrophage Infection. Cell Host Microbe 3, 323–330 (2008).

40. Luo, D. et al. Universal Stress Proteins: From Gene to Function. Int. J. Mol. Sci. 24, 4725 (2023).

41. De Maio, A. HEAT SHOCK PROTEINS: FACTS, THOUGHTS, AND DREAMS. Shock 11, 1 (1999).

42. Caño-Muñiz, S., Anthony, R., Niemann, S. & Alffenaar, J.-W. C. New Approaches and Therapeutic Options for Mycobacterium tuberculosis in a Dormant State. Clin. Microbiol. Rev. 31, 10.1128/cmr.00060-17 (2017).

43. Pai, M. P., Cottrell, M. L., Kashuba, A. D. M. & Bertino, J. S. 19 - Pharmacokinetics and Pharmacodynamics of Anti-infective Agents. in Mandell, Douglas, and Bennett’s Principles and Practice of Infectious Diseases (Eighth Edition) (eds. Bennett, J. E., Dolin, R. & Blaser, M. J.) 252–262.e2 (W.B. Saunders, Philadelphia, 2015). doi:10.1016/B978-1-4557-4801-3.00019-9.

44. Rustomjee, R. et al. Early Bactericidal Activity and Pharmacokinetics of the Diarylquinoline TMC207 in Treatment of Pulmonary Tuberculosis. Antimicrob. Agents Chemother. 52, 2831 (2008).

45. Wynn, E. A. Emergence of antibiotic-specific Mycobacterium tuberculosis phenotypes during prolonged treatment of mice. Antimicrob Agents Chemother (IN PRESS).

46. Mandell, Douglas & Bennett. Mandell, Douglas, and Bennett’s Principles and Practice of Infectious Diseases. https://www.elsevier-elibrary.com/product/mandell-douglas-bennetts-principles-practice-infectious-diseases-ebook (2014).

47. Vilchèze, C. et al. Commonalities of Mycobacterium tuberculosis Transcriptomes in Response to Defined Persisting Macrophage Stresses. Front. Immunol. 13, 909904 (2022).

48. Eagle, H. & Musselman, A. D. The Slow Recovery of Bacteria from the Toxic Effects of Penicillin. J. Bacteriol. 58, 475–490 (1949).

49. Srimani, J. K., Huang, S., Lopatkin, A. J. & You, L. Drug detoxification dynamics explain the postantibiotic effect. Mol. Syst. Biol. 13, 948 (2017).

50. Zhang, N. et al. Mechanistic Modeling of Mycobacterium tuberculosis Infection in Murine Models for Drug and Vaccine E=icacy Studies. Antimicrob. Agents Chemother. 64, 10.1128/aac.01727-19 (2020).

51. Dartois, V. Drug forgiveness and interpatient pharmacokinetic variability in tuberculosis. J. Infect. Dis. 204, 1827–1829 (2011).

52. Fox, W. S., Strydom, N., Imperial, M. Z., Jarlsberg, L. & Savic, R. M. Examining nonadherence in the treatment of tuberculosis: The patterns that lead to failure. Br. J. Clin. Pharmacol. 89, 1965–1977 (2023).

53. Li, R. Correlation between bactericidal activity and postantibiotic effect for five antibiotics with different mechanisms of action. J. Antimicrob. Chemother. 40, 39–45 (1997).

