## Supplemental Document 1 for "Physiologic recovery of *Mycobacterium tuberculosis* from drug injury: A molecular study of post antibiotic effect in mice"

### 1. Supplemental Methods

#### 1.1. Bacterial bioinformatics

After sequencing, bacterial reads were trimmed by Skewer v0.2.2[1] using an adapter sequence of 5'- CTGTCTCTTATACACATCT-3', a length of 50-175 bases, and an end quality of 20. Individual reads were reunited with their mate pair using PairFQ lite v0.17.0.[2] Paired-end reads were then aligned to the 2016 *Mtb* Erdman reference assembly using Bowtie2 v2.5.0[3] with default parameters. Mapped sequences were counted using HtSeq v1.0[4] with the default parameters.

The SEARCH-TB panel amplifies *Mtb* 3,568 genes. In this experiment, 12 genes had high incidences of 0 expression values, so our analysis included 3,552 genes.

#### 1.2. Table S1. Drug dosages

**Table S1**. Drugs and doses used in treatment with the HRZE regimen

| Drug | Dosage | Abbreviation |
|---|---|---|
| Isoniazid | 10m/kg | H |
| Rifampin | 10mg/kg | R |
| Pyrazinamide | 150mg/kg | Z |
| Ethambutol | 100mg/kc | E |

#### 1.3. Table S1. Curated gene categories

**Table S2**. Curated gene categories used for average expression analysis

Gene categories curated from the literature were used in category enrichment analysis. Along with the reference, the total number of genes from each source along with the number of genes from each source which were in the SEARCH-TB assay is given. Other gene sets used for enrichment analysis are available in Cole *et al.*, 1998.[5]

| **Category** | **Genes Total** | **Genes In Assay** | **Source** |
|---|---|---|---|
| ABC transporters | 72 | 68 | 6 |
| ABC transporters - Anion Transporters | 7 | 7 | 7 |
| ABC transporters - Metal Transporters | 5 | 5 | 7 |
| ABC transporters - Type I peptide and amino acids | 8 | 6 | 7 |
| ABC transporters - Type I phosphate | 8 | 8 | 7 |
| ABC transporters - Type I Sugar Importers | 12 | 12 | 7 |
| Alternative ribosomal proteins | 5 | 4 | 8 |
| Antigen 85 | 3 | 3 | 9 |
| Antitoxins | 76 | 72 | 10 |
| Arabinogalactan (AG) | 20 | 19 | 11 |
| Beta Oxidation | 18 | 17 | 12 |
| Cell wall synthesis | 40 | 40 | 13 |
| Cholesterol A and B ring degradation | 10 | 10 | 14 |
| Cholesterol C and D ring degradation | 5 | 5 | 14 |
| Cholesterol side chain degradation | 33 | 33 | 14 |
| Cutinase-Like Protein (CULP) | 7 | 7 | 15 |
| Cytochrome *bcc/aa3* supercomplex | 7 | 6 | 16 |
| Cytochrome *bd* oxidase | 4 | 3 | 16 |
| Diacyltrehalose (DAT) and Pentaacyltrehalose (PAT) | 5 | 5 | 17 |
| DNA replication and repair | 27 | 25 | 18 |
| DosR | 48 | 48 | 19 |

| | | | |
|---|---|---|---|
| Efflux Pumps and Transports | 26 | 25 | [20] |
| Enduring Hypoxic Response | 161 | 149 | [21] |
| Esterases (Lip family) | 22 | 20 | [15] |
| ESX1 | 19 | 18 | [22] |
| ESX2 | 12 | 12 | [22] |
| ESX3 | 11 | 9 | [22] |
| ESX4 | 7 | 7 | [22] |
| ESX5 | 15 | 11 | [22] |
| Fatty Acid Synthases I | 1 | 1 | [5] |
| Fatty Acid Synthases II | 10 | 9 | [23] |
| Fumarate reductase | 4 | 4 | [5] |
| Kas operon | 5 | 3 | [24] |
| kstR1 regulon | 74 | 70 | [25] |
| kstR2 regulon | 15 | 14 | [25] |
| LAM | 15 | 14 | [26] |
| LpqY-SugA-SugB-Sug trehalose transporter | 5 | 5 | [7] |
| Mce1 | 7 | 7 | [5] |
| Mce2 | 7 | 6 | [5] |
| Mce3 | 7 | 7 | [5] |
| Mce4 | 7 | 7 | [5] |
| mmpL | 14 | 14 | [27] |
| mmpS | 5 | 5 | [28] |
| Mycobactin Biogenesis | 10 | 10 | [29] |
| Mycolic Acid Modification | 12 | 12 | [30] |
| Mycolic Acid Transfer and Modification | 7 | 7 | [30] |
| NADH dehydrogenase type I | 14 | 12 | [31] |
| NADH dehydrogenase type II | 2 | 2 | [31] |
| Nitrate import and reductase | 6 | 8 | [5] |
| Oxidative Stress | 51 | 50 | [32] |
| Phthiocerol Dimycocerosate (PDIM) | 20 | 20 | [33] |
| Peptidoglycan (PG) | 34 | 32 | [34] |
| PG synthesis1- cytoplasmic steps | 7 | 7 | [34] |
| PG synthesis2- membrane-associated steps | 3 | 3 | [34] |

| | | | |
|---|---|---|---|
| Peptidoglycan (PG) Linking | 13 | 13 | [34] |
| PG synthesis4- remodeling and degradation | 2 | 2 | [34] |
| Peptidoglycan (PG) Transport and recycling | 3 | 3 | [34] |
| Phospholipases C | 4 | 4 | [35] |
| Primary Ribosomal Protein | 53 | 50 | [5] |
| response to acid stress | 11 | 9 | |
| Ribosomal hibernation | 4 | 4 | [36] |
| Sigma Factors | 13 | 12 | [6] |
| Stringent Response - Induced | 70 | 58 | [37] |
| Stringent Response - Repressed | 78 | 66 | [37] |
| Succinate Dehydrogenase Types I and II | 7 | 7 | [38] |
| Toxin-Antitoxin | 152 | 143 | [10] |
| Toxins | 76 | 74 | [10] |
| Transcription Factors | 198 | 187 | [6] |
| Trehalose | 5 | 5 | [39] |
| Triacylglycerol Synthases | 16 | 14 | [40] |
| UgpABCE glycerophosphocholine transporter | 4 | 4 | [7] |
| Universal Stress Proteins | 10 | 9 | [6] |
| WhiB-Like transcription factors | 7 | 7 | [41] |
| Zur regulon | 20 | 17 | [42] |

* Primary Ribosomal Proteins include ribosomal proteins identified by Cole *et al.* but exclude the four alternative ribosomal proteins

### 2. Supplemental Results

#### 2.1. Comparison of transcriptome at end of treatment compared to pre-treatment control

Treatment with HRZE for two and four weeks massively transformed the *Mtb* transcriptome, significantly altering expression of 2,497 (70.3%) and 2,617 (73.7%) genes relative to pre-treatment control, respectively.

**a** **b**

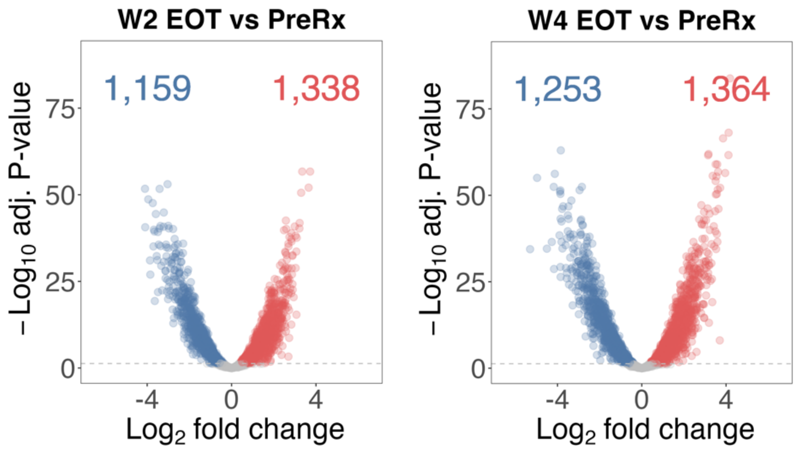

**Fig. S1. Comparison of transcriptome at end of treatment compared to pre-treatment control.** Volcano plot showing $\log_2$ fold changes and $-\log_{10}$ P-values of genes differentially expressed at the end of a two-week (a) or four-week (b) treatment compared to pre-treatment. Genes significantly down- and up-regulated relative to control (adj. $P < 0.05$) are shown in blue and red, respectively.

#### 2.2. Concordance of effect of treatment on expression with previous study

Gene expression changes after two and four weeks of treatment were compared to previous results published by Wynn *et al.*[43] The $\log_2$ fold change at the end of treatment compared to the pre-treatment (PreRx) control were determined for both studies and plotted against each other for two (**Fig. S2a**) and four (**Fig. S2b**) weeks of treatment. Expression changes of most genes were in the same direction, though $\log_2$ fold change in Wynn *et al.* tended to be smaller.

a b

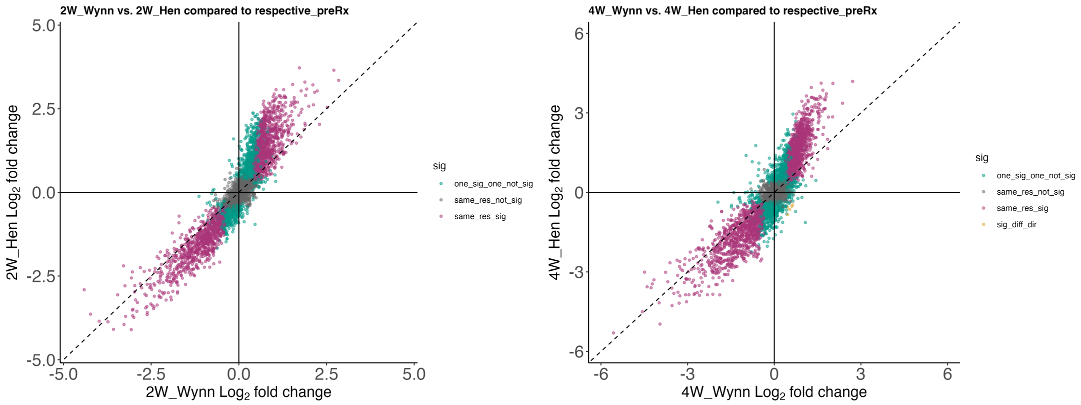

**Fig. S2. Concordance of effect of treatment on expression with previous study.** Comparison of two (A) or four (B) weeks of HRZE treatment versus control fold changes from this study versus Wynn *et al.* Purple shading indicates genes with concordant fold-change direction and significance between current study and Wynn *et al.* Green shading indicates genes that were significant in current or Wynn *et al.* results but not both. Gold shading indicates genes that were significant for both studies but in opposite directions. Gray shading indicates genes that were not significantly differentially expressed in either study.

#### 2.3. Effect of HRZE treatment length on gene expression

The $log_2$ fold change in gene expression after four weeks of HRZE compared to PreRx control were compared to the $log_2$ fold change in gene expression after two weeks of HRZE compared to PreRx control. Although the number of genes differentially expressed between two and four weeks was not large, the four-week transcriptome had larger $log_2$ fold changes than two weeks of treatment suggesting that the phenotype of *Mtb* treated with HRZE for four weeks is a more extreme version of the phenotype observed after two weeks of treatment (**Fig. S3**).

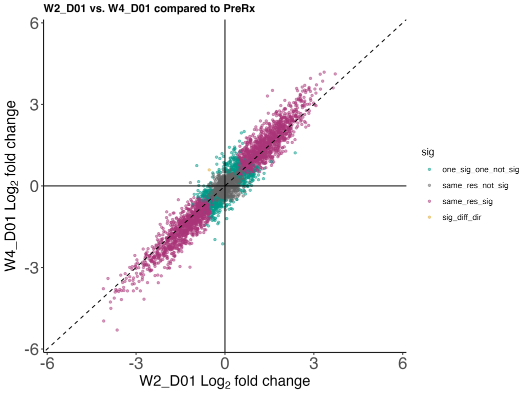

**Fig. S3. Effect of HRZE treatment length on gene expression**. Comparison of two versus four weeks of HRZE treatment versus control. Purple shading indicates genes with concordant fold-change direction and significance between two- and four-week treatment. Green shading indicates genes that were significant in two- or four-week results but not both. Gold shading indicates genes that were significant for both treatment groups but in opposite directions. Gray shading indicates genes that were not significantly differentially expressed in either treatment groups.

#### 2.4. Comparison of CFU and RS ratio to baseline

The median CFU and RS ratio of each time point during the PAE phase were compared to the baseline (day 0) with calculated adjusted P-value (padj).

**Table S3.** Recovery of CFU after 2 weeks of treatment

| Time | Median $Log_{10}$ CFU | padj (Comparing timepoint to day 1) |
|---|---|---|
| 1 | 5.86 | 1 |
| 5 | 6.09 | 0.80100176 |
| 7 | 5.93 | 0.91389221 |
| 11 | 5.77 | 0.80100176 |
| 14 | 5.85 | 0.80100176 |
| 21 | 5.81 | 0.80100176 |
| 28 | 6.13 | 0.90985585 |

**Table S4.** Recovery of CFU after 4 weeks of treatment

| Time | Median $Log_{10}$ CFU | padj (Comparing timepoint to day 1) |
|---|---|---|
| 1 | 5.41 | 1 |
| 4 | 5.22 | 0.89515405 |
| 7 | 5.10 | 0.58715587 |
| 11 | 4.78 | 0.30197165 |
| 14 | 4.61 | 0.30197165 |
| 21 | 5.26 | 1 |
| 28 | 5.40 | 1 |

**Table S5.** Recovery of RS ratio after 2 weeks of treatment

| Time | Median RS ratio | padj (Comparing timepoint to day 1) |
|---|---|---|
| 1 | 20.52 | 1 |
| 5 | 47.48 | 0.00590896 |
| 7 | 48.91 | 0.00239611 |
| 11 | 57.97 | 0.00011954 |
| 14 | 61.20 | 0.00139673 |

| | | |
|---|---|---|
| 21 | 69.67 | 0.00122635 |
| 28 | 78.57 | 0.00309677 |

**Table S6.** Recovery of RS ratio after 4 weeks of treatment

| Time | Median RS ratio | padj (Comparing timepoint to day 1) |
|---|---|---|
| 1 | 16.95 | 1 |
| 4 | 30.55 | 0.01542229 |
| 7 | 36.22 | 0.00456429 |
| 11 | 38.83 | 0.00456429 |
| 14 | 35.67 | 0.02057123 |
| 21 | 35.76 | 0.02109756 |
| 28 | 43.06 | 0.01623013 |

#### 2.5. Gene expression of gene categories related to central carbon metabolism

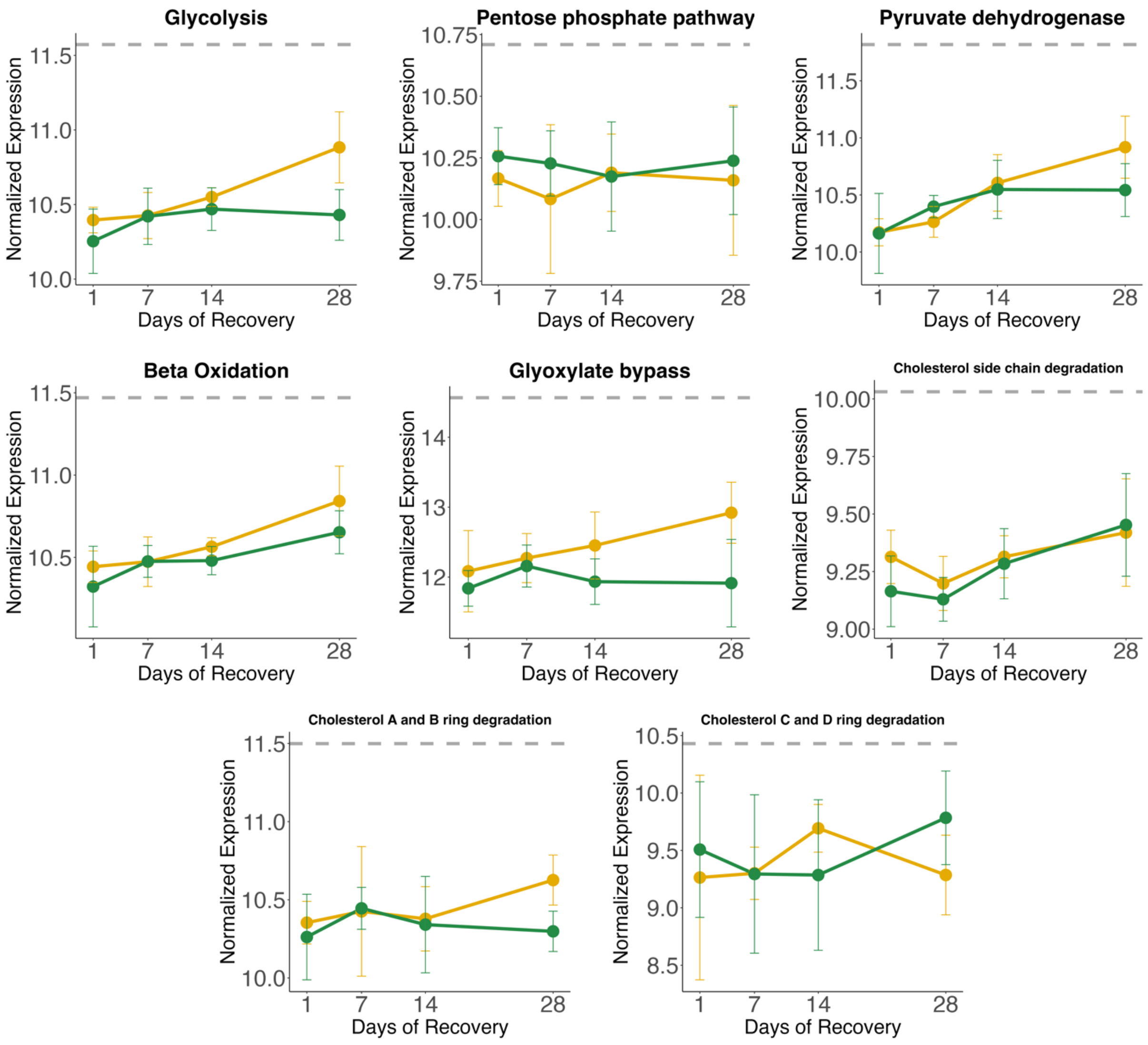

**Fig. S4. Gene expression of gene categories related to central carbon metabolism.** Normalized gene expression of select gene categories during treatment interruption following two (yellow) or four (green) weeks of treatment. Points indicate the average expression level and whiskers indicate 95% Confidence Interval between the five samples per time point. Gray line indicates average expression level of PreRx control.

#### 2.6. Gene expression of gene categories related to electron transport

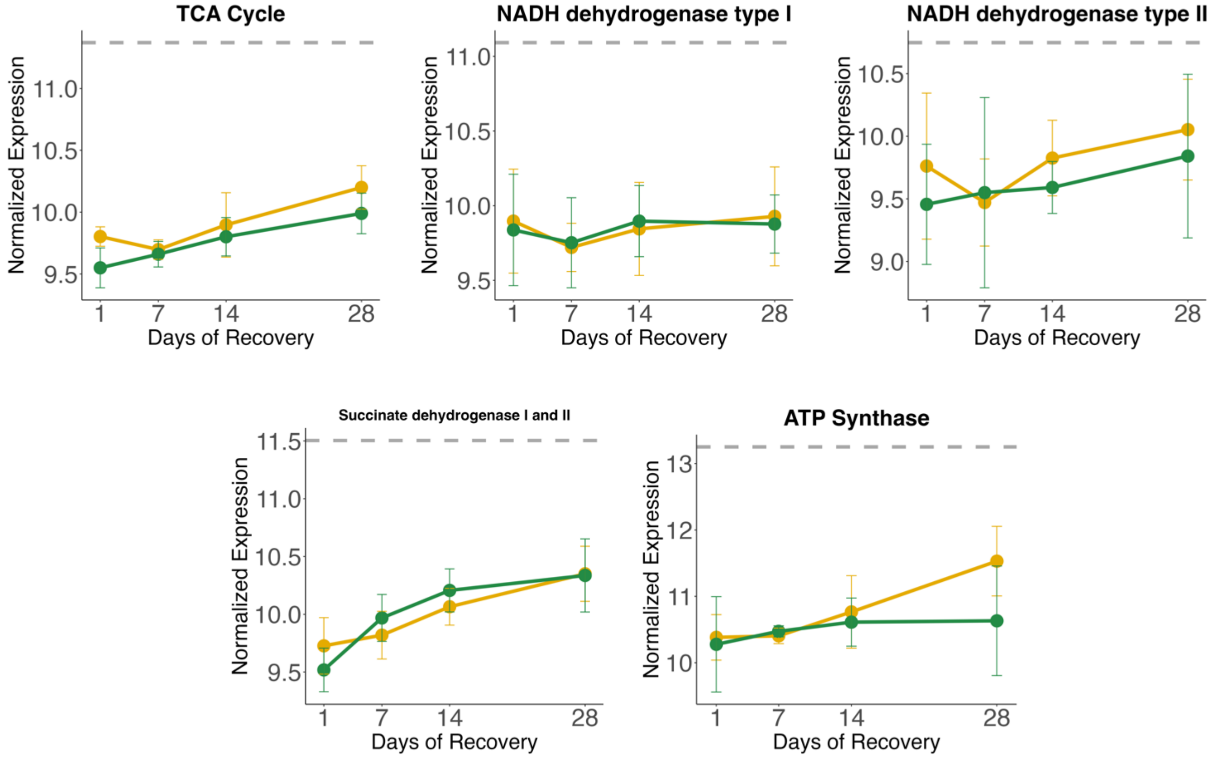

**Fig. S5. Gene expression of gene categories related to electron transport.** Normalized gene expression of select gene categories during treatment interruption following two (yellow) or four (green) weeks of treatment. Points indicate the average expression level and whiskers indicate 95% Confidence Interval between the five samples per time point. Gray line indicates average expression level of PreRx control.

#### 2.7. Gene expression of sigma factors

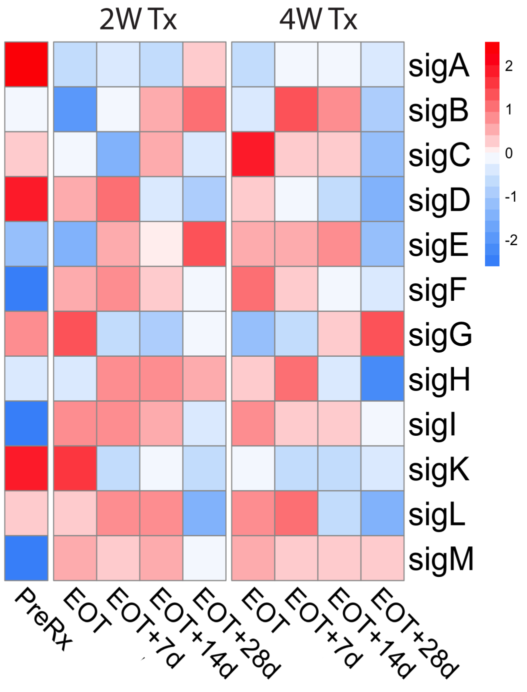

**Fig. S6. Gene expression of sigma factors.** Heatmap of normalized gene expression of sigma factor genes for the PreRx control, end of two and four weeks of treatment, and after 7, 14, and 28 days of post-antibiotic recovery for each treatment group.

#### 2.8. Comparison of transcriptome after recovery by treatment duration

The transcriptomes of *Mtb* treated for two and four weeks were compared following the 28-day post-antibiotic period. More genes were differentially expressed after the post-antibiotic period (N = 229) than at the end of treatment (N = 759), suggesting that *Mtb* recovered differently depending on treatment duration.

**a** **b**

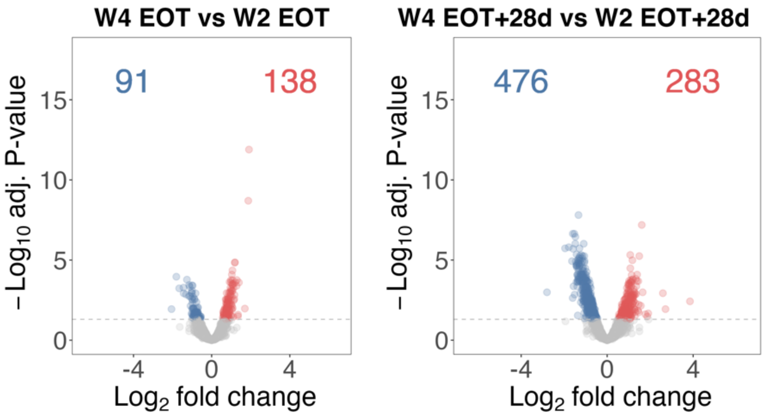

**Fig. S6. Comparison of transcriptome after recovery by treatment duration.** Volcano plot showing $\log_2$ fold changes and $-\log_{10}$ P-values of genes differentially expressed at the end of treatment (a) and at the end of post-antibiotic recovery period (b) following four versus two weeks of treatment. Genes significantly down- and up-regulated relative to control (adj. $P < 0.05$) are shown in blue and red, respectively.

### 3. References

1. Jiang, H., Lei, R., Ding, S.-W. & Zhu, S. Skewer: a fast and accurate adapter trimmer for next-generation sequencing paired-end reads. *BMC Bioinformatics* **15**, 182 (2014).
2. Staton, E. Pairfq. (2022).
3. Langmead, B., Trapnell, C., Pop, M. & Salzberg, S. L. Ultrafast and memory-efficient alignment of short DNA sequences to the human genome. *Genome Biol.* **10**, R25 (2009).
4. Putri, G. H., Anders, S., Pyl, P. T., Pimanda, J. E. & Zanini, F. Analysing high-throughput sequencing data in Python with HTSeq 2.0. *Bioinformatics* **38**, 2943–2945 (2022).
5. Cole, S. T. *et al.* Deciphering the biology of Mycobacterium tuberculosis from the complete genome sequence. *Nature* **396**, 190–190 (1998).
6. Lew, J. M., Kapopoulou, A., Jones, L. M. & Cole, S. T. TubercuList – 10 years after. *Tuberculosis* **91**, 1–7 (2011).
7. Soni, D. K., Dubey, S. K. & Bhatnagar, R. ATP-binding cassette (ABC) import systems of Mycobacterium tuberculosis: target for drug and vaccine development. *Emerg. Microbes Infect.* **9**, 207–220 (2020).
8. Kushwaha, A. K. & Bhushan, S. Unique structural features of the Mycobacterium ribosome. *Prog. Biophys. Mol. Biol.* **152**, 15–24 (2020).
9. Karbalaei Zadeh Babaki, M., Soleimanpour, S. & Rezaee, S. A. Antigen 85 complex as a powerful Mycobacterium tuberculosis immunogene: Biology, immune-pathogenicity, applications in diagnosis, and vaccine design. *Microb. Pathog.* **112**, 20–29 (2017).

10. Shao, Y. *et al.* TADB: a web-based resource for Type 2 toxin–antitoxin loci in bacteria and archaea. *Nucleic Acids Res.* **39**, D606–D611 (2011).

11. Abrahams, K. A. & Besra, G. S. Synthesis and recycling of the mycobacterial cell envelope. *Curr. Opin. Microbiol.* **60**, 58–65 (2021).

12. Schnappinger, D. *et al.* Transcriptional Adaptation of Mycobacterium tuberculosis within Macrophages: Insights into the Phagosomal Environment. *J. Exp. Med.* **198**, 693–704 (2003).

13. Kirksey, M. A. *et al.* Spontaneous phthiocerol dimycocerosate-deficient variants of Mycobacterium tuberculosis are susceptible to gamma interferon-mediated immunity. *Infect. Immun.* **79**, 2829–2838 (2011).

14. Pawełczyk, J. *et al.* Cholesterol-dependent transcriptome remodeling reveals new insight into the contribution of cholesterol to Mycobacterium tuberculosis pathogenesis. *Sci. Rep.* **11**, 12396 (2021).

15. Tallman, K. R., Levine, S. R. & Beatty, K. E. Small-Molecule Probes Reveal Esterases with Persistent Activity in Dormant and Reactivating Mycobacterium tuberculosis. *ACS Infect. Dis.* **2**, 936–944 (2016).

16. Lee, B. S., Sviriaeva, E. & Pethe, K. Targeting the cytochrome oxidases for drug development in mycobacteria. *Prog. Biophys. Mol. Biol.* **152**, 45–54 (2020).

17. Nobre, A., Alarico, S., Maranha, A., Mendes, V. & Empadinhas, N. The molecular biology of mycobacterial trehalose in the quest for advanced tuberculosis therapies. *Microbiology* **160**, 1547–1570 (2014).

18. Ditse, Z., Lamers, M. H. & Warner, D. F. DNA Replication in Mycobacterium tuberculosis. *Microbiol. Spectr.* **5**, (2017).

19. Voskuil, M. I. *et al.* Inhibition of Respiration by Nitric Oxide Induces a *Mycobacterium tuberculosis* Dormancy Program. *J. Exp. Med.* **198**, 705–713 (2003).

20. Remm, S., Earp, J. C., Dick, T., Dartois, V. & Seeger, M. A. Critical discussion on drug efflux in Mycobacterium tuberculosis. *FEMS Microbiol. Rev.* **46**, fuab050 (2021).

21. Rustad, T. R., Harrell, M. I., Liao, R. & Sherman, D. R. The Enduring Hypoxic Response of Mycobacterium tuberculosis. *PLOS ONE* **3**, e1502 (2008).

22. Gröschel, M. I., Sayes, F., Simeone, R., Majlessi, L. & Brosch, R. ESX secretion systems: mycobacterial evolution to counter host immunity. *Nat. Rev. Microbiol.* **14**, 677–691 (2016).

23. Duan, X., Xiang, X. & Xie, J. Crucial components of mycobacterium type II fatty acid biosynthesis (Fas-II) and their inhibitors. *FEMS Microbiol. Lett.* **360**, 87–99 (2014).

24. Slayden, R. A. & Barry, C. E. The role of KasA and KasB in the biosynthesis of meromycolic acids and isoniazid resistance in Mycobacterium tuberculosis. *Tuberc. Edinb. Scotl.* **82**, 149–160 (2002).

25. Wipperman, M. F., Sampson, N. S. & Thomas, S. T. Pathogen roid rage: cholesterol utilization by Mycobacterium tuberculosis. *Crit. Rev. Biochem. Mol. Biol.* **49**, 269–293 (2014).

26. Batt, S. M., Burke, C. E., Moorey, A. R. & Besra, G. S. Antibiotics and resistance: the two-sided coin of the mycobacterial cell wall. *Cell Surf. Amst. Neth.* **6**, 100044 (2020).

27. Domenech, P., Reed, M. B. & Barry, C. E. Contribution of the Mycobacterium tuberculosis MmpL protein family to virulence and drug resistance. *Infect. Immun.* **73**, 3492–3501 (2005).

28. Melly, G. & Purdy, G. E. MmpL Proteins in Physiology and Pathogenesis of M. tuberculosis. *Microorganisms* **7**, 70 (2019).

29. Quadri, L. E. N., Sello, J., Keating, T. A., Weinreb, P. H. & Walsh, C. T. Identification of a *Mycobacterium tuberculosis* gene cluster encoding the biosynthetic enzymes for assembly of the virulence-conferring siderophore mycobactin. *Chem. Biol.* **5**, 631–645 (1998).

30. Marrakchi, H., Lanéelle, M.-A. & Daffé, M. Mycolic Acids: Structures, Biosynthesis, and Beyond. *Chem. Biol.* **21**, 67–85 (2014).

31. Cook, G. M., Hards, K., Vilchèze, C., Hartman, T. & Berney, M. Energetics of Respiration and Oxidative Phosphorylation in Mycobacteria. *Microbiol. Spectr.* **2**, (2014).

32. Voskuil, M. I., Bartek, I., Visconti, K. & Schoolnik, G. K. The Response of Mycobacterium Tuberculosis to Reactive Oxygen and Nitrogen Species. *Front. Microbiol.* **2**, (2011).

33. Rens, C., Chao, J. D., Sexton, D. L., Tocheva, E. I. & Av-Gay, Y. Roles for phthiocerol dimycocerosate lipids in Mycobacterium tuberculosis pathogenesis. *Microbiol. Read. Engl.* **167**, (2021).

34. Maitra, A. *et al.* Cell wall peptidoglycan in Mycobacterium tuberculosis: An Achilles' heel for the TB-causing pathogen. *FEMS Microbiol. Rev.* **43**, 548–575 (2019).

35. Raynaud, C. *et al.* Phospholipases C are involved in the virulence of Mycobacterium tuberculosis. *Mol. Microbiol.* **45**, 203–217 (2002).

36. Li, Y., Corro, J. H., Palmer, C. D. & Ojha, A. K. Progression from remodeling to hibernation of ribosomes in zinc-starved mycobacteria. *Proc. Natl. Acad. Sci.* **117**, 19528–19537 (2020).

37. Dahl, J. L. *et al.* The role of RelMtb-mediated adaptation to stationary phase in long-term persistence of Mycobacterium tuberculosis in mice. *Proc. Natl. Acad. Sci. U. S. A.* **100**, 10026–10031 (2003).

38. Hartman, T. *et al.* Succinate dehydrogenase is the regulator of respiration in Mycobacterium tuberculosis. *PLoS Pathog.* **10**, e1004510 (2014).

39. Thanna, S. & Sucheck, S. J. Targeting the trehalose utilization pathways of Mycobacterium tuberculosis. *MedChemComm* **7**, 69–85 (2016).

40. Maurya, R. K., Bharti, S. & Krishnan, M. Y. Triacylglycerols: Fuelling the Hibernating Mycobacterium tuberculosis. *Front. Cell. Infect. Microbiol.* **8**, 450 (2018).

41. Wan, T. *et al.* Structural insights into the functional divergence of WhiB-like proteins in *Mycobacterium tuberculosis*. *Mol. Cell* **81**, 2887-2900.e5 (2021).

42. Dow, A. *et al.* Zinc limitation triggers anticipatory adaptations in Mycobacterium tuberculosis. *PLoS Pathog.* **17**, e1009570 (2021).

43. Wynn, E. A. *et al.* Transcriptional adaptation of Mycobacterium tuberculosis that survives prolonged multi-drug treatment in mice. *mBio* **14**, e02363-23 (2023).
